# Genomic diversity and functional potential of facultative bacterial symbionts across scale insects

**DOI:** 10.1101/2025.10.27.684769

**Authors:** Pradeep Palanichamy, Jinyeong Choi, Arno Hagenbeek, Filip Husnik

## Abstract

Microbial symbionts play pivotal roles in the physiology, ecology, and evolution of insects. While obligate symbionts have been extensively characterized in insects feeding on nutritionally poor diets, the diversity and functional roles of facultative bacterial symbionts remain largely unexplored in many insect lineages, including scale insects (Hemiptera: Coccomorpha). Here, we present a genome-resolved metagenomic and comparative genomic analysis of facultative bacterial symbionts across 120 scale insect metagenomes from 20 different families. Our analyses reveal rich and taxonomically diverse facultative symbiont communities, dominated by the Pseudomonadota, such as *Wolbachia, Rickettsia, Arsenophonus*, and *Sodalis*. Genomic features reveal substantial variation in genome sizes, coding densities, metabolic potentials, and host-interaction genes among symbiont clades, indicating lineage-specific lifestyles and host interactions. Interestingly, alphaproteobacterial and gammaproteobacterial symbionts mostly co-occur within their respective hosts. Gammaproteobacterial symbionts exhibit broader metabolic repertoires and potential defense capabilities via the APSE phage toxin cassettes, while alphaproteobacterial symbionts retain reduced metabolic capabilities, Type IV secretion systems, and reproductive manipulation genes (*cifAB* and *wmk*). We propose how the symbiont genes related to nutritional provisioning, defensive symbiosis, and reproductive manipulation may influence the biology and evolution of scale insects. Our results provide the first comprehensive genomic overview of facultative bacterial symbionts in scale insects, revealing their evolutionary dynamics and putative functions.

## Introduction

Symbiotic interactions between insects and microorganisms have played and continue to play a significant role in the evolutionary success of many insect lineages^1,2^. From the insect host’s perspective, facultative symbionts (also known as secondary or accessory) are not essential for the insect’s survival or reproduction but can confer a wide range of benefits or costs, including protection against natural enemies, increased thermal tolerance, enhanced nutrition, and sex ratio distortion^3–7^. Depending on the symbiont lineage and insect host, facultative symbionts can be both intracellular and extracellular in various tissues, including hemolymph, gut, salivary glands, and reproductive organs^3,6,7^. Facultative symbiont titers are also contextdependent, and their infection frequencies can vary highly across populations, species, and seasons, often depending on environmental conditions and transmission fidelity^5,8^. While maternally inherited, horizontal transmission through parasitoids, host plants, or mating also occurs and plays a crucial role in their spread across unrelated insect lineages^3,6,9^.

Facultative symbionts represent a genomic intermediate between free-living bacteria and obligate mutualists, displaying signatures of genome reduction such as pseudogenization, reduced metabolic capacity, and accumulation of mobile genetic elements^10–13^. Despite these features, they often retain relatively large genomes and greater functional plasticity compared to obligate symbionts^10,14^. Notably, their genomes frequently harbor a diverse array of mobile elements such as transposons, integrative and conjugative elements (ICEs), prophages, and plasmids, which facilitate horizontal gene transfer and promote genomic rearrangements^15–17^.

Two major drivers of facultative symbiont evolution are secretion systems and viruses. Protein secretion systems are key mediators of prokaryotic-eukaryotic interactions, enabling symbionts to manipulate host physiology, evade immune responses, and establish intracellular niches–specialized environments within host cells that provide protection and access to host-derived nutrients^18,19^. Facultative bacterial symbionts often retain diverse Type III (T3SS), Type IV (T4SS), and Type VI secretion systems^10,20^. For instance, the T3SS in *Sodalis* may facilitate the initial colonization of host cells, while the T4SS in *Wolbachia* is implicated in effector protein delivery and reproductive manipulation^10,21^. The retention and functional diversification of these secretion systems underscore their potential role in facilitating facultative symbiosis and mediating symbi-ont-host communication^10,13^.

Bacteriophages are increasingly recognized as major players in the ecology and evolution of facultative symbionts^22^, acting as vectors for the horizontal transfer of genes encoding toxins or effector proteins that shape host-symbiont dynamics^10,23^. One of the best-characterized examples is the APSE (*Acyrthosiphon pisum* Secondary Endosymbiont) phage, which carries toxin cassettes such as Shiga-like toxins, cytolethal-distending toxins (*cdtB*), and YD-repeat toxins that confer protection against parasitoid wasps in aphids^24,25^. On the other hand, WO phages in *Wolbachia* have been implicated in reproductive manipulation strategies like cytoplasmic incompatibility and male killing ^26,27^. WO phages possess eukaryotic association modules (EAMs) and may carry genes associated with toxin-antitoxin systems, which contribute to *Wolbachia*’s ability to persist in host populations and affect host biology^28^.

Scale insects (Hemiptera: Coccomorpha) comprise a highly diverse group of sap-feeding insects, with over 8,500 described species^29^. They exhibit extreme morphological and ecological adaptations and occupy a wide range of ecological niches on diverse plants^30^. Like other plant sap-feeding hemipterans, scale insects rely on microbial partners to supplement their nutritionally imbalanced diets^31–34^. While the role of obligate bacterial symbionts in providing these insects with essential amino acids and B vitamins is well-studied, the identity, diversity, and functional significance of facultative bacterial symbionts in scale insects remain largely unknown^35^. Facultative bacterial symbionts, such as *Fritschea, Cardinium, Spiroplasma, Enterobacter*-like, *Wolbachia, Rickettsia, Arsenophonus, Sodalis*, and *Symbiopectobacterium*, have been previously identified in scale insects but have been rarely studied in detail^32,33,36–46^. Most current knowledge about facultative symbioses of hemipteran insects thus derives from studies in aphids, whiteflies, and psyllids, where symbionts such as *Hamiltonella, Serratia, Wolbachia, Arsenophonus, Rickettsia*, and *Cardinium* have been shown to affect host reproduction, defense, stress tolerance, and metabolic flexibility^47–50^. It is expected that similar symbiont dynamics may exist in scale insects. However, it remains to be tested, as scale insect biology in many ways deviates from other hemipterans.

Here, we present the diversity and comparative genomic analysis of facultative bacterial symbionts associated with scale insects. We analyzed metagenomes of 120 scale insect species across 20 different families collected from diverse host plants and habitats. Using genome-resolved metagenomes, we provide an in-depth phylogenomic and metabolic functional profiling of this understudied group. Our results reveal a diversity of facultative bacterial symbionts and illuminate their functional repertoires. These findings highlight scale insects as a rich but largely unexplored reservoir of insect symbionts, advancing our understanding of how facultative symbioses influence insect biology.

## Results

### Diversity and phylogenomic relationships of facultative bacterial symbionts across scale insects

From 120 scale insect samples, we identified 72 putative facultative bacterial symbionts spanning five bacterial phyla and ten families, including Mycoplasmatota (Entomoplasmatales), Actinomycetota (Micrococcales), Chlamydiota (Parachlamydiales), Bacteroidota (Cytophagales), and Pseudomonadota (Rhodospirillales, Sphingomonadales, Hyphomicrobiales, Rickettsiales, Pseudomonadales, Orbales, and Enterobacterales) (Fig. 1). At the genus level, *Wolbachia, Sodalis, Arsenophonus*, and *Rickettsia* were predominant, with *Wolbachia* found across nine scale insect families and *Rickettsia* associated with three scale insect families. *Sodalis* and *Arsenophonus* were each identified across four scale insect families (Fig. 1). To resolve their evolutionary relationships, we constructed a comprehensive phylogenomic tree based on 99 conserved singlecopy BUSCO genes from 58 metagenome-assembled genomes (MAGs), encompassing a broad diversity of symbionts (Fig. 2A). Most symbionts clustered within well-established genera, while lineages such as *Fritschea, Bombella*-like, *Gilliamella*-like, and *Hemipteriphilus* formed monophyletic groups distinct from their closely related counterparts (Fig. 2A). Phylogenomic analysis of *Wolbachia* using 232 BUSCO genes revealed four supergroups within scale insects, with five strains each in supergroups A and B, two in D (or a novel supergroup), and one in E (Fig. *2B)*. Similarly, *Rickettsia* phylogeny based on 91 BUSCO genes showed all five-scale insect-associated *Rickettsia* cluster within the Belli group (Fig. 2C). *Arsenophonus* phylogeny reconstructed using 169 BUSCO genes revealed that most strains fall within the Triatominarum clade, except *Arsenophonus* FASP, which formed a sister group to the Apicola clade (Fig. 2D). *Sodalis* phylogeny based on 257 BUSCO genes revealed two distinct clades among *Sodalis* symbionts in scale insects, with four strains in Clade A and four in Clade B (Fig. 2E). Across these four genera (*Wolbachia, Rickettsia, Arsenophonus, Sodalis*), average nucleotide identity (ANI) based clustering patterns were highly congruent with respective phylogenomic trees, further validating the robustness of the trees (Sup Fig. 1A, Fig. 1B, Fig. 1C, Fig. 1D). Facultative symbionts from Alphaproteobacteria and Gammaproteobacteria mostly co-occur together within the same scale insect host, especially in mealybugs, giant scales, and a few minor families (Supplementary Table ST6). These phylogenetic analyses have provided us with a groundwork for analyzing the symbiont genome features.

**Figure 1:**
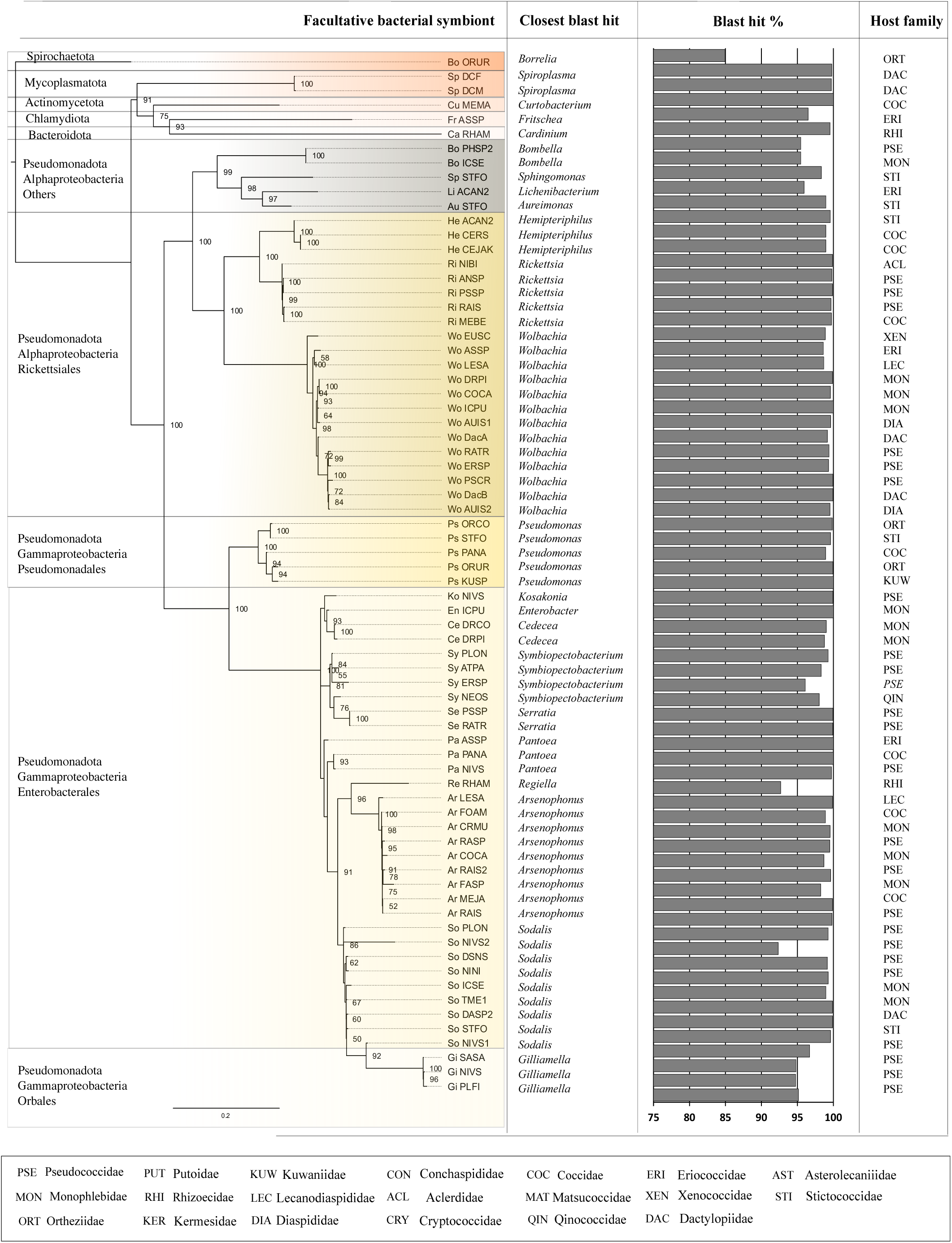
Maximum-likelihood tree from nearly complete sequences of the 16S rRNA gene of bacterial symbionts from scale insects. The tree was generated using IQtree under the TIM3+F+I+G4 model. The host is indicated in the three-letter code representing the family. The closest hit represents the best BLASTN hit of the 16S rRNA gene sequences against the 16S rRNA NCBI database. The bar plot represents the percentage identity of the best BLASTN hit. The bootstrap values below 50 are not shown. We note that due to poor phylogenetic signal in the 16S rRNA gene, the relationship between genera within Enterobacterales is largely unresolved.

**Figure 2:**
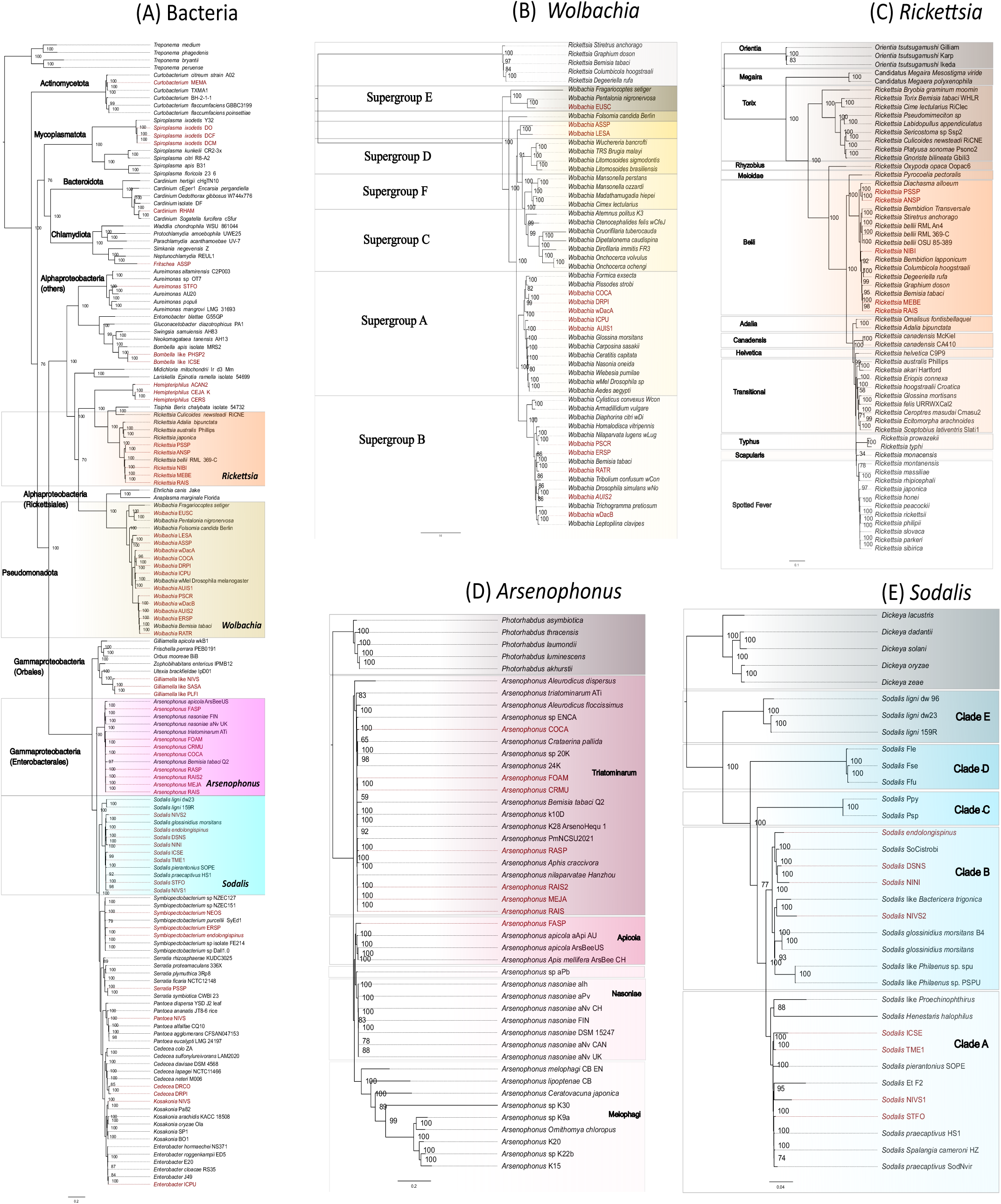
Maximum-likelihood phylogenies of diverse bacterial symbionts associated with scale insects. (A) Tree constructed from 99 singlecopy BUSCO genes (Bacteria) representing diverse bacterial phyla including Actinomycetota, Mycoplasmatota, Bacteroidota, Chlamydiota, and Pseudomonadota, using the LG+R9 model in IQ-TREE. (B) *Wolbachia* phylogeny based on 232 single-copy BUSCO genes (Rickettsiales) under the HIVb+F+R6 model. (C) *Rickettsia* phylogeny based on 91 single-copy BUSCO genes (Rickettsiales) under the JTT+F+R5 model. (D) *Ar-senophonus* phylogeny based on 169 single-copy BUSCO genes (Enterobacterales) under the cpREV+F+R5 model. (E) *Sodalis* phylogeny based on 257 single-copy BUSCO genes (Enterobacterales) under the JTT+R4 model. In all trees, symbionts from scale insects are highlighted in red, and most branches are supported by bootstrap values greater than 95.

### Genomic features of facultative bacterial symbionts from scale insects

A comprehensive analysis of genomic features was conducted for facultative bacterial symbionts encompassing 58 genomes across 17 genera. The BUSCO completeness of these genomes ranged from 67% to 99% (Fig. 3) (Supplementary Table). Their genome sizes and GC contents varied significantly, ranging from 1.07 to 6.76 Mbp and from 24.2 to 71.8 GC (Fig. 3). Their coding density ranged from 56% to 90%.

**Figure 3:**
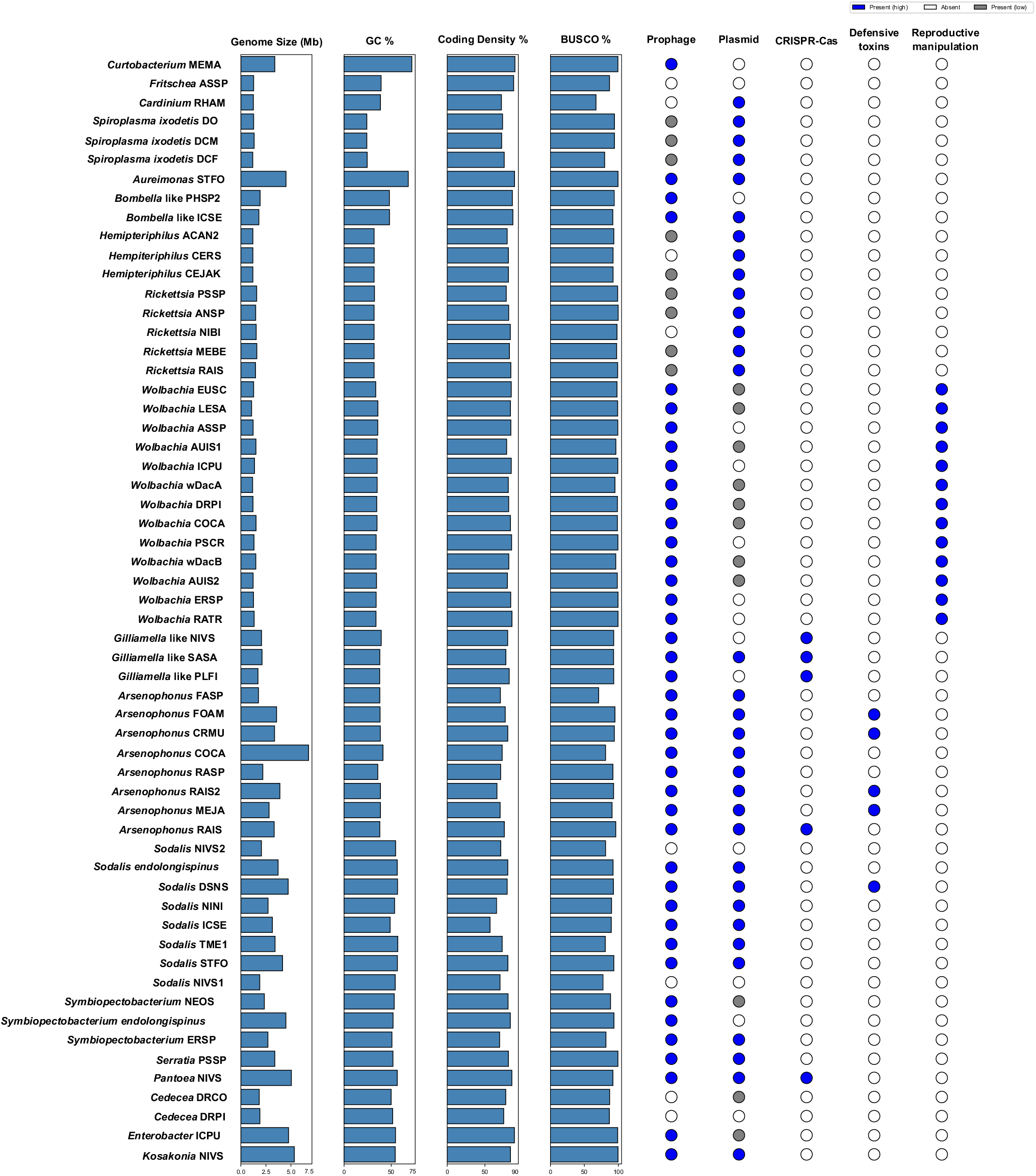
Genome features of bacterial symbionts from scale insects. The horizontal bar plot represents genomic features as follows in order from left to right: genome size (Mb), GC content (%), coding density (%), BUSCO scores (%) against the lineage-specific database. The circle plots indicate the presence and absence of prophage, plasmids, CRISPR-Cas system, defense toxins, and reproductive manipulation (filled in blue: present with higher confidence, in grey: present with lower confidence, in white: not present).

Plasmids and prophage elements were present in several genomes (Fig.3), categorized as either high-confidence or lowconfidence predictions. Most of the prophages identified within Pseudomonadota symbionts belonged to the phylum Uroviricota (Caudoviricetes). Prophages identified in *Wolbachia* genomes belong to the WO phage group (Supplementary Table ST4). Prophages identified in *Arsenophonus* and some *Sodalis* genomes belong to the APSE-like phage group (Supplementary Table ST4). Finally, the presence of the CRISPR-Cas defense system was identified in some of the symbiont genomes (Fig. 3). Genomic analyses of the facultative symbionts revealed the presence of candidate genes linked to reproductive manipulation and defensive symbiosis. In *Wolbachia* genomes, we identified key reproductive manipulation genes, including *cifA, cifB*, and *wmk* (Supplementary Fig. 5A, 5B, 5C), which are associated with cytoplasmic incompatibility and male killing. These genes were located within the WO prophage regions, consistent with their known mobile genetic context. In contrast, the genomes of *Arsenophonus* and *Sodalis* harbored putative defensive symbiosis genes, such as *cdtB*, Shiga-like toxins, and YD-repeat toxins, many of which were embedded within the APSE-like phage regions (Supplementary Fig. 4A, 4B, 4C).

**Figure 4:**
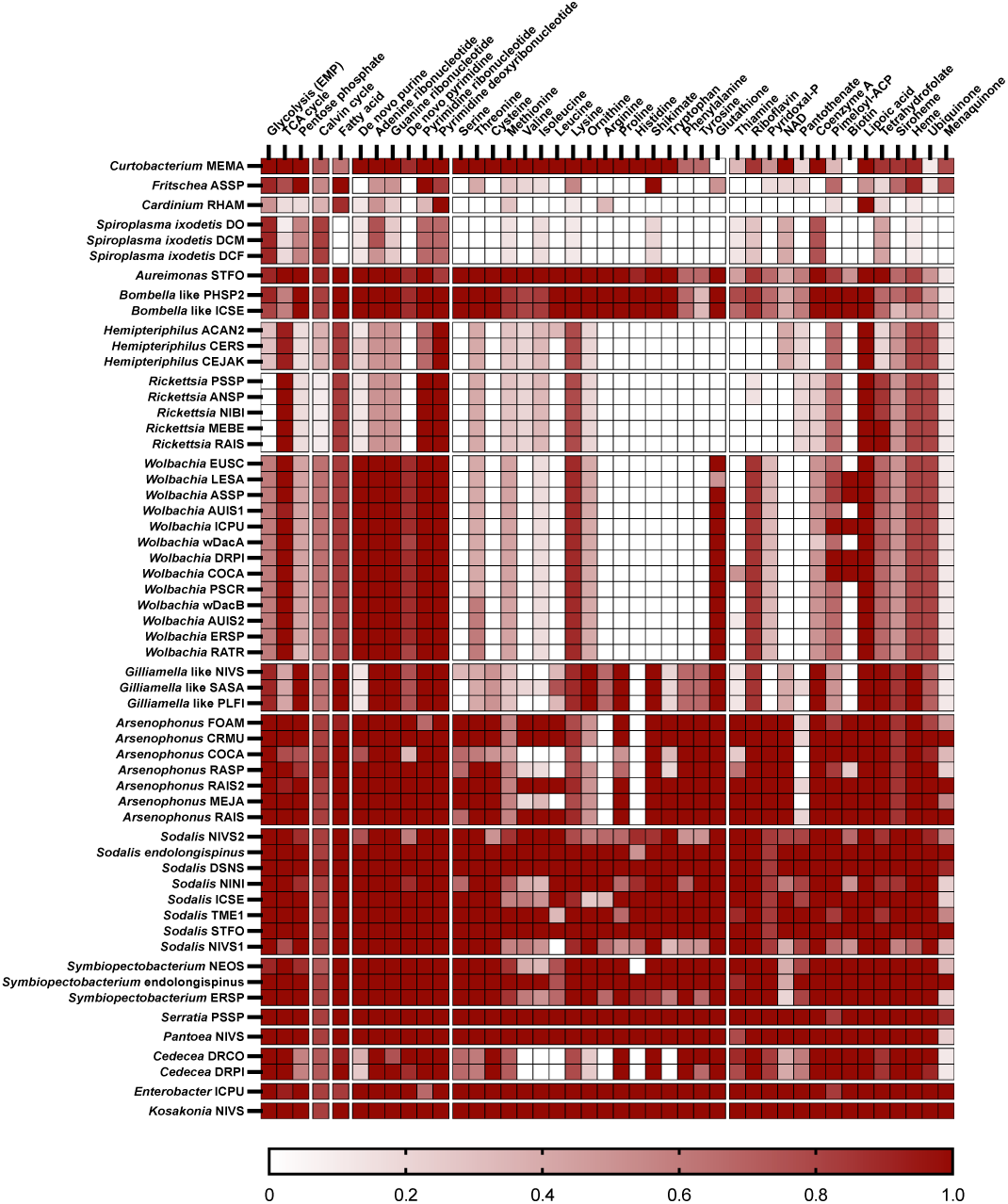
Metabolic capabilities of bacterial symbionts in scale insects. The heatmap indicates genomic completeness of major metabolic pathways related to carbohydrate metabolism, energy metabolism, lipid metabolism, nucleotide metabolism, amino acid metabolism, and metabolism of cofactors and vitamins.

**Figure 5:**
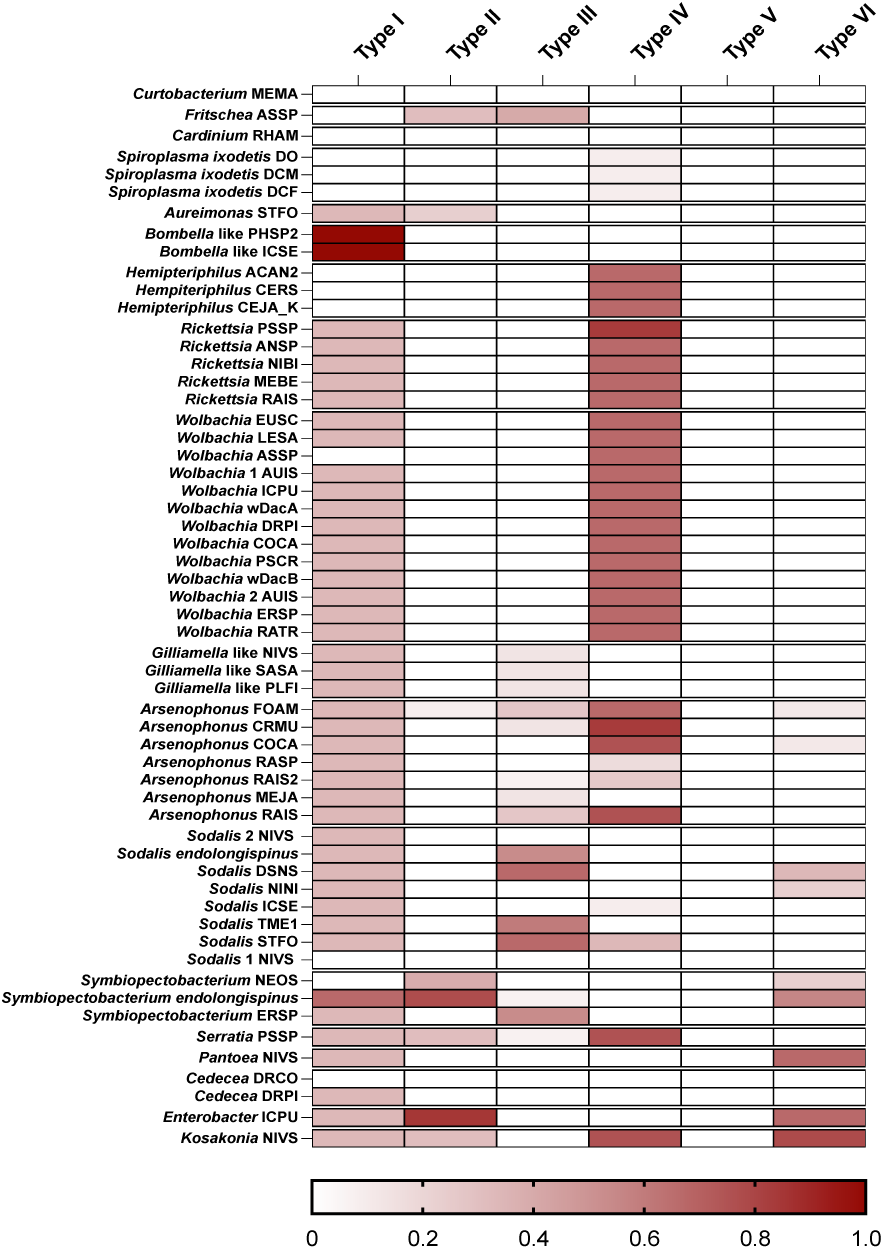
**Heatmap showing the completeness of bacterial type secretion systems**, including TypeI, Type II, Type II, Type IV, Type V, and Type VI systems, across the analyzed genomes. Completeness for each secretion system was calculated as the proportion of genes present in a genome relative to the total number of genes required for that secretion system.

### Functional potential of facultative bacterial symbionts

#### Carbohydrate and energy metabolism

The completeness of central metabolic pathways varied markedly among the facultative bacterial symbionts, reflecting distinct degrees of metabolic autonomy and host dependence (Fig.4). A fully complete glycolytic pathway, indicative of robust carbohydrate metabolism, was retained by *Curtobacterium* and several gammaproteobacterial symbionts. In contrast, only partial glycolytic capacity was detected in *Cardinium, Wolbachia*, and *Hemipteriphilus*, while glycolysis was absent in *Rickettsia*, consistent with their reduced genomes and obligate intracellular lifestyle. The tricarboxylic acid (TCA) cycle was fully complete in most metabolically capable taxa, including *Curtobacterium* and most of the alphaproteobacterial and gammaproteobacterial symbionts. Whereas a partially complete TCA cycle was observed in *Fritschea, Cardinium, Spiroplasma*, and *Bombella*-like and *Gilliamella*-like symbionts, suggesting varying degrees of respiratory capacity. The pentose phosphate pathway, was complete in a broad range of symbionts but reduced in *Cardinium, Spiroplasma, Cedecea*, and Rickettsiales symbionts that likely rely on host-derived intermediates to compensate for missing functions (Fig. 4).

Consistent with their variable respiratory potential, the distribution of genes encoding electron transport chain (ETC) components and ATP synthase complexes also differed across symbionts. Both ETC and ATP synthase genes were present in *Curtobacterium, Fritschea, Aureimonas, Bombella-like, Rickettsia, Wolbachia, Gilliamella-like, Arsenophonus, Sodalis, Symbiopectobacterium, Serratia, Pantoea, Cedecea, Enterobacter*, and *Kosakonia*, indicating the capacity for oxidative phosphorylation. In contrast, *Cardinium* and *Spiroplasma* lacked genes for the ETC, whereas *Hemipteriphilus* lacked ATP synthase subunits. Notably, *Fritschea* and several other symbionts retained only partial ETC components, further supporting the trend of stepwise metabolic reduction among certain lineages (Supplementary Table ST7).

#### Lipid metabolism

Lipid biosynthesis pathway reconstructions revealed variable completeness, reflecting the potential host dependence of some of the facultative symbionts (Fig.4). The fatty acid biosynthesis pathway was complete in *Fritschea, Aureimonas, Bombella*-like, and most of the gammaproteobacterial symbionts, suggesting these bacteria retain the capacity for *de novo* membrane lipid synthesis. A nearly complete or partially complete pathway was detected in *Cardinium, Curtobacterium*, and Rickettsiales symbionts. *Spiroplasma* lacked all key genes for fatty acid biosynthesis, indicating a strong reliance on host-derived lipids (Fig. 4).

#### Nucleotide metabolism

Similarly, the purine biosynthesis pathway was complete in *Curtobacterium* and most of the alphaproteobacterial and gammaproteobacterial symbionts, while partial gene loss was observed in *Fritschea, Cardinium, Spiroplasma, Hemipteriphilus, Rickettsia, Cedecea*, and *Gilliamella*-like symbionts. In contrast, the pyrimidine biosynthesis pathway displayed more intricate patterns of gene retention and loss. Most of the gammaproteobacterial symbionts retained a complete set of genes, although partial loss was noted in *Gilliamella*-like symbionts. Among Alphaproteobacteria, *Wolbachia, Aureimonas*, and *Bombella*-like symbionts possess most, or all, of the genes required for pyrimidine biosynthesis. In contrast, *Fritschea, Cardinium, Spiroplasma, Hemipteriphilus*, and *Rickettsia* pathways exhibited substantial reductions, including the complete absence of the *de novo* pathway and variable retention of genes for ribonucleotide and deoxyribonucleotide biosynthesis (Fig. 4).

#### Amino acid metabolism

A comparative analysis of amino acid biosynthesis pathways revealed a striking lineage-specific variation, reflecting differential metabolic capabilities and potential host dependency (Fig. 4). The actinobacterial symbiont *Curtobacterium* retained a nearly complete repertoire for amino acid biosynthesis, lacking only the phenylalanine and tyrosine pathways. In contrast, *Fritschea* and *Cardinium* exhibited widespread loss of amino acid biosynthesis genes, retaining only partial shikimate pathway genes in *Fritschea. Spiroplasma* displayed a limited capacity, with only partial methionine, isoleucine, lysine, and shikimate pathways. Among Alphaproteobacteria, *Aureimonas* and *Bombella*-like symbionts retained most amino acid biosynthesis pathways but shared partial losses in phenylalanine and tyrosine biosynthesis, with *Bombella*-like symbionts also showing losses in methionine, valine, and isoleucine pathways.

Rickettsiales symbionts showed broad losses in multiple pathways, accompanied by partial reductions in threonine, methionine, isoleucine, and ornithine biosynthesis. Notably, valine biosynthesis was absent in *Wolbachia* and partially degraded in *Hemipteriphilus* and *Rickettsia*. While glutathione biosynthesis was lacking in *Hemipteriphilus* and *Rickettsia*, it remains intact in most *Wolbachia*. Interestingly, lysine biosynthesis remained conserved across all Rickettsiales symbionts. Gammaproteobacterial symbionts generally maintained broader biosynthetic capacity, although degradation of biosynthesis pathways for serine, threonine, cysteine, methionine, valine, isoleucine, leucine, ornithine, proline, tryptophan, phenylalanine, tyrosine, arginine, and histidine was observed across *Gilliamella*-like symbionts, *Arsenophonus, Sodalis, Symbiopectobacterium*, and *Cedecea*. Notably, *Arsenophonus* showed complete loss of both arginine and histidine biosynthesis pathways, while *Gilliamella*-like symbionts and *Cedecea* had completely lost the biosynthetic pathways for valine, isoleucine, leucine, arginine, histidine, and tryptophan. In contrast, lysine, shikimate, and glutathione biosynthesis remained largely intact. *Serratia, Pantoea, Enterobacter*, and *Kosakonia* maintained a complete suite of genes for all amino acid biosynthesis pathways, underscoring their metabolic independence when contrasted to the more reduced symbiont lineages (Fig. 4).

#### B-vitamin and cofactor biosynthesis

Facultative bacterial symbionts of scale insects exhibit substantial variation in the retention and degradation of their B-vitamin and cofactor biosynthetic pathways across taxonomic groups. *Curtobacterium* retains complete or nearly complete pathways for riboflavin, NAD, coenzyme A, lipoic acid, tetrahydrofolate, siroheme, heme, and menaquinone, but shows partial gene loss in thiamine, pyridoxal-P, pantothenate, pimeloyl-ACP, and ubiquinone, with biotin biosynthesis absent. *Fritschea* retains most genes for heme and menaquinone, but lacks those for thiamine, riboflavin, coenzyme A, and biotin. *Cardinium* retains only the lipoic acid pathway, while *Spiroplasma* preserves most genes for coenzyme A but shows degradation in thiamine, riboflavin, NAD, tetrahydrofolate, and heme. Among alphaproteobacterial symbionts, riboflavin biosynthesis is absent in *Hemipteriphilus* and *Rickettsia*. Coenzyme A is fully retained in *Aureimonas* and *Bombella*-like symbionts, partially degraded in *Wolbachia* and *Rickettsia*, and absent in *Hemipteriphilus*. Biotin biosynthesis is absent in *Hemipteriphilus, Rickettsia*, and some *Wolbachia* strains, reduced in *Aureimonas*, and complete in *Bombella*-like symbionts and other *Wolbachia* strains. Tetrahydrofolate and pimeloyl-ACP pathways show partial gene loss across multiple taxa, while lipoic acid biosynthesis remains intact in all. The heme and ubiquinone pathways are largely conserved, especially among *Rickettsiales*. Gammaproteobacterial symbionts generally retain complete pathways for riboflavin, coenzyme A, lipoic acid, tetrahydrofolate, and heme (except in *Sodalis* NIVS1), but exhibit varying levels of loss in other pathways. For example, thiamine biosynthesis is partially reduced in *Gilliamella*-like symbionts and some *Arsenophonus* strains, and biotin biosynthesis is degraded in *Arsenophonus* RASP and some *Sodalis* strains, but remains complete in most others (Fig. 4).

### Secretion systems

Multiple scale insect bacterial symbionts (*Aureimonas, Rickettsia, Wolbachia, Gilliamella*-like symbiont, *Arsenophonus, Sodalis, Symbiopectobacterium, Serratia, Pantoea, Cedecea, Enterobacter*, and *Kosakonia*) exhibit a loss of genes related to the Type I secretion system or show evidence of T1SS disruption. Notably, genes associated with the Type I secretion system are present in *Bombella*-like symbionts. The Type II secretion system (T2SS) has undergone reduction in *Fritschea, Aureimonas, Arsenophonus* FOAM, *Symbiopectobacterium* NEOS, as well as in *Serratia* and *Kosakonia*. In contrast, *Symbiopectobacterium endolongispinus* and *Enterobacter* ICPU retain most of the genes associated with the T2SS. The Type III secretion system (T3SS) is reduced in several genera, including *Fritschea, Gilliamella*-like, *Arsenophonus, Sodalis, Symbiopectobacterium*, and *Serratia*. The majority of T3SS-related genes are present in *Sodalis* strains (*S. endolongispinus*, DSNS, TME1, STFO). Genes required for the Type IV secretion system (T4SS) are predominantly present in *Hemipteriphilus, Rickettsia, Wolbachia*, and most *Arsenophonus, Serratia*, and *Kosakonia* strains. However, the T4SS is absent in *Spiroplasma*, and also missing from certain *Arsenophonus* and *Sodalis* strains, suggesting either an ancestral absence, lineage specific acquisition or secondary gene loss depending on lineage history. In contrast, the Type VI secretion system (T6SS) is consistently present in *Kosakonia, Enterobacter*, and *Pantoea*, which retain most of the core genes required for its function. By comparison, T6SS genes are lacking in *Symbiopectobacterium* and are variably absent among *Sodalis* and *Arsenophonus* strains, indicating independent losses or lineage-specific absence rather than a uniform reduction across these genera (Fig. 5).

## Discussion

### Pseudomonadota are the most prevalent facultative bacterial symbionts in scale insects

Pseudomonadota are the dominant facultative symbionts in scale insects, accounting for more than 90% of our data. Pseudomonadota, including *Cedecea*, which is closely related to *Enterobacter*^36,39,43^, *Wolbachia*^42,44^, *Rickettsia*^45^, *Arsenophonus*^40^, *Sodalis*^32,33,37^, *Symbiopectobacterium*^32,37^, *Gilliamella*-like^51^, and *Sphingomonas*^41^, have been previously reported as facultative symbionts of scale insects. In addition, several additional Pseudomonadota symbionts were newly identified by this study. Most of these symbionts have been previously reported from other insects, such as *Aureimonas*^52,53^, *Hemipteriphilus*^54,55^, *Serratia*^48^, *Pantoea*^56^, *Enterobacter*^57^, *Kosakonia*^58^, *Bombella*-like^59^, and *Pseudomonas*^60,61^. Facultative symbionts from other phyla, such as *Fritschea*^62^, *Cardinium*^38^, and *Spiroplasma*^46^, have also been previously reported in scale insects. *Curtobacterium*, a rare actinobacterial symbiont, was previously identified in other insects^63^.

The diversity of facultative bacterial symbionts in scale insects is similar to that in whiteflies^47^, aphids^48^, and treehoppers^59^, likely because they share the same ecological niche on plants. Similar to other insect hosts where *Wolbachia* dominates symbiotic associations^64^, both previous studies and our analyses demonstrate that *Wolbachia* is widely distributed across scale insects from a wide range of families^44^ (Fig. 1). This pattern suggests a potentially significant role of *Wolbachia* in scale insect biology. Insect-associated *Wolbachia* symbionts are predominantly classified into supergroups A and B, a pattern also observed in *Wolbachia* from scale insects (Fig. 2B)^64^. Similarly, *Rickettsia* symbionts in hemipteran insects, including those in scale insects (Fig. 2C), are largely affiliated with the Belli group ^65^. *Arsenophonus* endosymbionts from hemipterans, including scale insects (Fig. 2D), are primarily placed within the Triatominarum clade^13^. In contrast, *Sodalis* symbionts associated with hemipterans tend to cluster into either the *Sodalis praecaptivus* or *Sodalis glossinidius* clades, both of which include symbionts from scale insects (Fig. 2E)^12^. Co-occurrence of facultative bacterial symbionts from Alphaproteobacteria and Gammaproteobacteria was observed in scale insects, as previously reported from other hemipteran insects such as whiteflies^66,67^ and aphids ^7,68,69.^

### Gammaproteobacterial symbionts encode diverse metabolic pathways

Gammaproteobacterial facultative symbionts in scale insects possess diverse metabolic pathways, particularly related to the biosynthesis of essential amino acids and B vitamins. This metabolic versatility suggests that they may play a crucial role in providing nutrients to their scale insect hosts. One notable category was the enrichment of B vitamin biosynthesis genes (thiamine, riboflavin, pyridoxal-P, NAD, pantothenate, biotin, and folate), in agreement with previous studies from other insects ^37,67,70–72^. The completeness of these pathways in *Serratia, Pantoea, Enterobacter*, and *Kosakonia* suggests that these bacteria have recently transitioned from a free-living to an insect-associated lifestyle^6^. While diverse amino acid and cofactor pathways were specifically retained across the gammaproteobacterial symbionts, the biosynthesis pathways for lysine, glutathione, coenzyme A, lipoic acid, heme, and ubiquinone remained consistently conserved across all analyzed strains. This conservation implies that these compounds are primarily essential for core cellular functions, redox homeostasis, and energy metabolism in the bacterial symbionts, but they could also support the insect growth and fitness^73–77^.

### Alphaproteobacterial symbionts lack essential genes for nutrient biosynthesis

Alphaproteobacterial symbionts, within the order Rickettsiales (*Rickettsia, Wolbachia*, and *Hemipteriphilus*) are metabolically dependent on their insect hosts or co-occurring microbial symbionts. The notable loss of glycolysis-related genes is observed in *Rickettsia*, along with partial degradation of glycolysis genes in *Hemipteriphilus* and *Wolbachia*. This pattern is consistent with previous work on *Rickettsia*, which is well-known to rely on host-derived sugars and ATP^65^. Most amino acid biosynthesis pathways were absent in Rickettsiales, including serine, cysteine, valine, leucine, arginine, proline, histidine, shikimate, tryptophan, phenylalanine, and tyrosine. Partial loss was also observed in threonine, methionine, isoleucine, and ornithine biosynthesis, suggesting an ongoing transition toward greater metabolic dependency^42,78^.

B-vitamin biosynthesis was notably reduced, with thiamine, pyridoxal-P, NAD, pantothenate, and coenzyme A biosynthesis genes lacking across most Rickettsiales symbionts. Folate biosynthesis showed a gradual degradation across symbionts within this group. Despite these extensive losses, some pathways remain stable. Lysine biosynthesis was conserved across Rickettsiales^65,79^, likely due to the role of meso-diaminopimelic acid in peptidoglycan biosynthesis^80^, while glutathione biosynthesis was consistently maintained in *Wolbachia*, possibly aiding in oxidative stress resistance^81^. The presence of tetrahydrofolate biosynthesis genes in some *Rickettsia* strains (MEBE and RAIS) also suggests that they retain a degree of metabolic independence (Fig. 4)^82^. Interestingly, biotin biosynthesis genes were found in five *Wolbachia* symbionts of scale insects (LESA, ASSP, ICPU, DRPI, and COCA) (Fig. 4). These genes were acquired via horizontal gene transfer^83,84^ and, depending on the insect clade and diet, can also support the host^85^. Riboflavin biosynthesis genes appeared to be stable in *Wolbachia* symbionts within scale insects (Fig. 4), further highlighting lineage-specific metabolic retention patterns^86^. *Wolbachia* may supplement riboflavin and biotin to offset the fitness cost of its presence and influence the reproduction of some scale insects ^87^. On the other hand, the presence of horizontally acquired biotin and riboflavin biosynthesis genes on some scale insect genomes^31,32^ might eliminate the need for symbiont-derived biotin and riboflavin.

### *Arsenophonus* and *Sodalis* are potential defensive symbionts of scale insects

In addition to their potential metabolic contribution, our analysis identified several candidate genes associated with defensive symbiosis within four *Arsenophonus* and one *Sodalis* strains. These include *cdtB*, Shiga-like toxin genes and YD-repeat toxin genes, many of which were located in APSE-like prophage elements. These proteins are known to play protective roles in other insects by defending the host against parasitoids^24,88,89^. For instance, homologous toxin genes in *Hamiltonella defensa* have been shown to be essential for conferring resistance to parasitoid wasps, with their phage-borne nature being an important factor. We note that all the scale insect species from which we detected the potentially defensive symbionts are known to be hosts of parasitoid wasps^90–92^. This finding highlights the potential of these facultative symbionts to act as defensive mutualists. Under the selection pressure of scale insect natural enemies, *Arsenophonus* and *Sodalis* may be more than mere passengers and contribute to their host’s survival by producing toxins.

### *Wolbachia* is likely an active reproductive manipulator of scale insects

Our results provide genomic evidence that many *Wolbachia* strains associated with scale insects harbor genes linked to reproductive manipulation. Notably, we identified *cifA* and *cifB*, which are associated with cytoplasmic incompatibility, as well as *wmk*, a candidate male-killing gene. These genes were localized within WO phage regions, consistent with previous studies indicating that reproductive manipulation genes are often encoded within mobile genetic elements^27,93^. This aligns with earlier findings demonstrating the role of phage elements in shaping host-bacterial symbiont interactions^24,27,94–98^.

Scale insects are renowned for their remarkably diverse array of genetic systems, including diplodiploidy, haplodiploidy, paternal genome elimination (PGE), arrhenotoky, hermaphroditism, and various forms of parthenogenesis^99,100^. Transitions between these systems appear to have occurred multiple times, and evolutionary theory suggests that inter-genomic conflict (for example, between hosts and symbionts) may drive the emergence or maintenance of these diverse systems^99,100^. The presence of *Wolbachia* strains carrying genes related to cytoplasmic incompatibility, male killing, feminization, and parthenogenesis thus raises the possibility that reproductive manipulators may contribute not only to modifying reproduction but also to facilitating or selecting for transitions among the extremely diverse genetic systems found in scale insects. Together, these findings suggest that *Wolbachia* may act as an active reproductive manipulator and also potentially contribute to the evolution and diversity of reproductive and genetic systems.

### Type IV secretion system (T4SS) is essential for Rickettsiales symbionts of scale insects

The Type IV secretion system (T4SS) is highly conserved across Rickettsiales symbionts of scale insects, including *Wolbachia, Rickettsia*, and *Hemipteriphilus*. All analyzed genomes encode the core T4SS components (*virB3, virB6, virB8, virB9, virB10, virB4, virB11*, and *virD4*) indicating a stable and potentially functional T4SS across these lineages. Similar conservation has been reported in other intracellular Rickettsiales, where T4SS-mediated effector secretion plays a key role in host-microbe interactions^21,101,102^. In *Wolbachia, vir* gene clusters are highly conserved and transcriptionally active, particularly in the ovaries of infected arthropods, implicating the T4SS in the delivery of effector proteins that mediate reproductive manipulation^21^. In *Rickettsia*, the T4SS contributes to intracellular survival and immune evasion by secreting various effector proteins within host cells^103^.

T4SS is also a key mediator involved in horizontal gene transfer to other bacteria and eukaryotic cells^104^. Genomic fragments from *Wolbachia* and other Rickettsiales have been found to be integrated and transcriptionally active in various arthropod and nematode hosts, highlighting the potential of the T4SS to mediate gene transfer across domains^105^. Additionally, T4SS systems are known to facilitate DNA uptake, toxin secretion, and virulence factor delivery in diverse bacterial lineages^104,106^. We hypothesize that Rickettsiales symbionts of scale insects also employ the T4SS for intracellular persistence, host reproductive manipulation, and mediating HGT between co-infecting symbionts (and accidentally, with their eukaryotic hosts).

### Reproductive manipulation, nutrition, and defense are the putative roles of facultative symbionts across scale insects

In this manuscript, we identified a diverse array of bacterial symbionts associated with scale insects, including *Wolbachia, Rickettsia, Arsenophonus, Sodalis, Spiroplasma, Cardinium, Serratia, Symbiopectobacterium, Pantoea*, and several other bacterial lineages. Genomic and functional potential analyses suggest that these symbionts may contribute to a wide range of roles, pending experimental validation. By drawing parallels to their functional roles in other insect hosts, we hypothesize that they are involved in reproductive manipulation, nutrition, and defense.

For example, several of these symbionts are well-known reproductive manipulators. *Wolbachia* is widely recognized for its capacity to manipulate host reproduction through mechanisms such as cytoplasmic incompatibility, male killing, and parthenogenesis^64,107,108^. *Cardinium* induces similar effects, including cytoplasmic incompatibility, feminization, and parthenogenesis ^109–111^. *Spiroplasma* is implicated in male killing in various insects^112^, and reproductive manipulations by *Rickettsia* and *Arsenophonus* have also been documented^113–115^, suggesting that these bacteria may also influence reproductive dynamics in scale insects. Since we identified marker genes of reproductive manipulation in scale-insect-associated symbionts, our data support that they are likely involved in sex distortion phenotypes.

Beyond reproductive manipulation, many of the identified symbionts are known to contribute to host nutrition in a contextdependent manner. For instance, *Wolbachia* can provide biotin and riboflavin in nutrient-limited environments^87^. *Sodalis, Symbiopectobacterium*, and *Arsenophonus* often serve as co-obligate nutritional mutualists, including in plant sap-feeding insects^32,37,67^. *Serratia* has been reported to enhance fatty acid metabolism and supply essential nutrients to insect hosts^116,117^. Based on our comparative genomics data, these symbionts may similarly contribute to the nutritional and physiological needs of scale insects that feed not only on plant phloem, but also on parenchyma cells, xylem, and fungal hyphae^30^.

Another important functional axis revealed by our work is the defensive potential of the studied symbionts. Both *Spiroplasma* and *Rickettsia* confer protection against parasitoids and pathogens in other insects^118–120^. In whiteflies, *Rickettsia* strains very closely related to those detected by us in scale insects (the Belli group) were shown to confer resistance to entomopathogens such as *Pseudomonas syringae* and *Cordyceps javanica*^120,121^.

Finally, *Arsenophonus* and *Sodalis* strains related to those found in scale insects harbor APSE-like phages, which may provide protection against parasitoids^88,89^.

## Conclusion

Our results revealed the presence of genes for reproductive manipulation, nutrient biosynthesis, and defense across most facultative symbionts detected in scale insects. However, we acknowledge that these data should be interpreted with caution. The precise functional role of individual symbionts within specific hosts is likely to be dependent on variables such as parasitoid pressure, host plant diet, and environmental conditions. Since our primary focus was on the initial screening of a large diversity of both symbionts and hosts, this manuscript lays the groundwork for future experimental work. Nevertheless, the observed genomic features, coupled with evidence from other insects, support reproductive manipulation, nutrient provisioning, defense, and environmental adaptation as the core facultative symbiont capabilities. To pinpoint the context-dependent roles of specific facultative symbionts, we advocate for detailed experimental validation as the necessary next step.

## Acknowledgements

FH was supported by the JSPS KAKENHI grant (23K14256) and the HFSP Early Career Grant (RGEC29/2024). JC was supported by the National Research Foundation of Korea (NRF) grant funded by the Korean government (MSIT; 2021R1A6A3A03038909), and a JSPS KAKENHI grant (20939772). We also acknowledge the Scientific Computing (SCDA), and Sequencing (SQC) sections of the Okinawa Institute of Science and Technology for their great support.

## Author contributions

PP led the study, conducted DNA extraction, data analyses, prepared all figures, and drafted the manuscript. FH designed the study, supervised the analyses, and revised the manuscript. JC performed DNA extraction, data analysis and revised the manuscript. AH developed and optimized the bioinformatic pipelines used in this study. All authors edited the manuscript.

## Data availability

The genome assemblies of facultative bacterial symbionts and Illumina raw reads are available under the NCBI BioProject PRJNA1129605.

## Materials and Methods

### Scale insect sampling, DNA extraction, and genome sequencing

Scale insect samples were collected by our unit members and collaborators. All the insect samples were stored in 99% ethanol at −20°C. A few individuals from each species were selected, and wax secretions were carefully removed under the Olympus SZX16 dissecting microscope using fine forceps. Individuals showing signs of parasitoid infection were excluded during this process. Surface-sterilized insects were subsequently homogenized in liquid nitrogen using a mortar and pestle. Genomic DNA extractions were carried out with the MasterPure Complete DNA purification kit (Epicenter), according to the manufacturer’s instructions. Genomic DNA quantity and quality were assessed using a microvolume UV-Vis spectrophotometer (NanoDrop One; Thermo Fisher Scientific) and a Qubit 4 fluorometer (Invitrogen). PCR-free Illumina libraries were prepared using the NEBNext Ultra II protocol (NEB), multiplexed, and sequenced using the Illumina NovaSeq6000 and NovaSeqX platforms at the Okinawa Institute of Science and Technology and Pennsylvania State University. Genomic DNA extracted from three scale insect species was prepared for long-read sequencing using both Oxford Nanopore Technologies (ONT) and Pacific Biosciences (PacBio) platforms. For ONT sequencing libraries were prepared with the Native Barcoding Kit 96 V14 (SQK-NBD114.96) and sequenced on an R10.4.1 PromethION Flow Cell using the PromethION platform. For PacBio sequencing, libraries were prepared using the SMRTbell Express Template Prep Kit 2.0 and sequenced on the PacBio Revio system at the Okinawa Institute of Science and Technology.

### Metagenome screening for symbionts

All commands and codes are available in this GitHub repository https://github.com/ECBSU/ECBSU_manuscripts_code/tree/main/Facultative_bacterial_symbiont_scale_insect_manuscript. In addition to newly generated data, raw metagenomic reads from scale insects (n = 14) were retrieved from the Sequence Read Archive (SRA) of the National Center for Biotechnology Information (NCBI). In parallel, eight previously published bacterial symbiont genomes associated with scale insects were obtained from NCBI GenBank. These genomes were assessed for completeness using BUSCO v5.1.3^122^, employing lineage-specific bacterial databases as described below. Additionally, closely related bacterial genomes were retrieved from NCBI GenBank to serve as reference genomes for phylogenomic analyses. The approximate taxonomic affiliations and relative abundance of bacterial symbionts were estimated from small subunit rRNA read abundance in metagenomic data, analyzed using phyloFlash v3.4^123^.

### Genome assemblies

The quality of the raw Illumina reads was evaluated using FastQC v0.11.7^124^. Low-quality reads and adapter sequences were removed using fastp v0.20.0^125^. Read error correction, de novo assembly using multiple k-mers, and mismatch/short indel correction were carried out using the SPAdes assembler, v3.15^126^. Taxonomic classification of the assembled scaffolds was performed by conducting a megablast search (NCBI-BLAST v2.11.0) against the NCBI nucleotide database (March 2023). Bacterial scaffolds were retrieved based on the taxon-annotated GC-coverage plots generated by Blobtools v1.1.^127^. The draft symbiont genome assemblies were subsequently polished using Pilon v1.24^128^.

We further analyzed four long-read metagenomic datasets derived from scale insects. Of these, two datasets were generated using Oxford Nanopore PromethION sequencing, and two were obtained from PacBio HiFi sequencing. Quality control of the raw long reads was performed using tools from the NanoPack suite^129^, including NanoQC v0.10.0 (for basic quality assessment), NanoStat v1.6.0 (for statistical summaries), and NanoPlot v1.44.1 (for visualization of read quality and length distributions). Adapter sequences were trimmed using FastpLong v0.3.0^125^, and Filtlong v0.2.1 (https://github.com/rrwick/Filtlong) was employed for quality-based read filtering. The filtered long-read datasets from both Oxford Nanopore and PacBio platforms were assembled using Flye v2.9^130^ with two iterations. Taxonomic classification of the assembled scaffolds was performed as described above. Bacterial genomes assembled from the Oxford Nanopore datasets were polished using Medaka v2.1 (https://github.com/nanoporetech/medaka), while assemblies from PacBio HiFi reads were polished using NextPolish2 v0.2.1^131^. To further correct mismatches, insertions, and deletions, the Medaka and NextPolish2 corrected assemblies were refined with Pilon v1.2.4^128^ using corresponding Illumina short-read data from scale insect hosts.

### Identification and classifications of obligate and facultative bacterial symbionts

Scale insects are typically colonized by two distinct types of endosymbiotic bacteria: obligate symbionts, which are tightly integrated and essential to the host biology, and facultative symbionts, which are more variable in their occurrence and function. We classified symbionts into obligate or facultative categories based on multiple criteria adapted from established criteria in the literature ^10,12,132,133^. Our classification relied on host specificity, prevalence patterns, phylogenetic association with the host, genome characteristics, and functional genomic content.

Obligate symbionts of scale insects typically exhibit strict host specificity, being confined to a particular host species or insect family, and are consistently present across all individuals/population/species/etc. within the same clade (with some exceptions if recently replaced). Phylogenetic analyses indicate that obligate symbionts co-diversify with their hosts, reflecting a long-term coevolutionary relationship^36,134^. Genomes of obligate bacterial symbionts typically exhibit extensive genome reduction <1 Mbp (though recent replacements exceed this threshold), low GC content, and gene repertoires limited primarily to essential genetic machineries and nutritional functions. Mobile genetic elements such as prophages, plasmids, and insertion sequences are typically absent or rare, and protein secretion systems, including T3SS, T4SS, or T6SS, are usually not detected.

Currently characterized obligate symbionts of scale insects include *Tremblaya spp*. and *Sodalis*-allied bacteria in Pseudococcidae^31,32^, *Bacteroidetes* and *Ophiocordyceps* spp. in Rastrococcinae^35^, *Ophiocordyceps* spp. in Coccidae^135^, *Walczuchella* sp. in Monophlebidae^36^, *Brownia* sp. in Rhizoecidae^136^, *Sodalis* sp. in Putoidae^137^, *Uzinura* sp. in Diaspididae^34^, and *Dactylopiibacterium* sp. in Dactylopiidae^138^. Some families, such as Matsucoccidae, appear to lack obligate symbionts^35^. Our metagenomic data support the widespread occurrence of diverse Bacteroidetes, Gammaproteobacteria, and *Ophiocordyceps* spp. as obligate symbionts across scale insect families (Choi et al., unpublished).

In contrast, facultative symbionts in scale insects are not consistently present across different individuals, populations, or species within the same family. These symbionts can be acquired or lost in response to environmental pressures and are often horizontally transmitted. They are often more resistant to insect antimicrobial peptides or remain undetected by the insect immune response due to their variable outer membrane proteins. Facultative symbionts typically have larger genome sizes (>1 Mbps), more variable GC content (often higher than that of obligate symbionts), and a more complex gene repertoire that includes functions related to nutrition, defense, reproductive manipulation, stress tolerance, and host interactions. Genomes of facultative symbionts are rich in mobile genetic elements such as plasmids, transposons, and prophages, and commonly encode one or more secretion systems (e.g., T3SS, T4SS, T6SS).

### Genome annotations and analyses

Symbiont genomes were annotated with Prokka v.1.14.6 (--compliant --rfam -- evalue 1e^-05^)^139^. We assessed the KEGG completion score with a custom pipeline (https://github.com/ECBSU/genomics-scripts/tree/main/Functional-annotation/KEGGstand_in_house_KEGG_annotation). Each bacterial genome was annotated using eggNOG-mapper v2^140^ to generate KEGG annotation. The genes required to complete each KEGG module were then assessed. The script parses the KEGG module definitions to find each possible combination of genes that complete the module, taking into account gene redundancies and non-essential genes. For each genome, the completion of the modules was calculated by comparing each gene combination against the gene presence/absence given by the EggNOG annotation, with the fraction of genes present in the most complete combination taken as the module completion. Average Nucleotide Identity (ANI) between all-vs-all bacterial genomes was calculated by fastANI v1.34 (https://github.com/ParBLiSS/FastANI). Hierarchical clustering of the ANI matrix was performed using the UPGMA method implemented in SciPy. The heatmap was generated using an ANI clustered matrix (https://github.com/moshi4/ANIclustermap). Genome completeness was evaluated using BUSCO v5.1.3^122^, employing lineage-specific bacterial databases (alphaproteobacteria_odb10, entomoplasmatales_odb10, rhodospirillales_odb10, bacteria_odb10, chlamydiae_odb10, gammaproteobacteria_odb10, rickettsiales_odb10, cytophagales_odb10, micrococcales_odb10, bacteroidetes-chlorobi_group_odb10, mycoplasmatales_odb10, bacteroidetes_odb10, enterobacterales_odb10, pseudomonadales_odb10) to ensure accurate identification of single-copy orthologous genes. Mobile genetic elements, such as prophage and plasmids, were predicted using geNomad v1.11^141^. Additionally, CRISPR-Cas systems were identified and classified using CRISPRCasTyper v1.8^142^.

### Phylogenetic and phylogenomic analyses

For phylogenetic analyses, 16S rRNA gene sequences were extracted from both raw metagenomic reads and assembled contigs using phyloFlash v3.4^123^ and aligned using MAFFT v7^143^. The alignments were trimmed using trimAL v1.4.1^144^ with the -automated1 option, and a maximum-likelihood tree was reconstructed using IQ-Tree v1.6.12^145^. The best-fitting substitution models were determined using ModelFinder^146^, and branch support was assessed with 1,000 ultrafast bootstrap replicates (UFBoot). For phylogenomic analyses, five separate phylogenomic trees were generated based on single-copy orthologous genes identified using BUSCO v5.1.3^122^ from genome assemblies of scale insect-associated bacterial symbionts and their close relatives (retrieved from NCBI GenBank). Three BUSCO lineage-specific databases were used: Bacteria_odb10 for all bacteria, Enterobacterales_odb10 for *Arsenophonus* and *Sodalis*, and Rickettsiales_odb10 for *Wolbachia* and *Rickettsia*. Protein sequences of orthologous genes were aligned using MAFFT v7^143^, trimmed with trimAL v1.4.1 (-automated1)^144^, and concatenated using a custom Python script (https://github.com/ECBSU/genomics-scripts/tree/main/multigene-phylogenomy/Busco_gene_multigene_tree). Maximum likelihood phylogenomic trees were constructed using IQ-TREE v.1.6.9^145^, with substitution models selected by ModelFinder^146^ and node support evaluated using 1000 ultrafast bootstrap replicates. All trees were visualized using FigTree v1.4.4^147^ and further refined in Inkscape.

### Identification of reproductive manipulation and defensive symbiosis genes

To identify the presence of reproductive manipulation and defensive symbiosis-associated genes, we first compiled a curated dataset of reference genes. This included cytoplasmic incompatibility factor genes (*cifA, cifB*), male-killing gene (*wmk*), and parthenogenesis factor genes (*pifA, pifB*) from various Wolbachia strains, as well as known defensive symbiosis genes such as *cdtB*, shiga-like toxin, and YD-repeat domain-containing genes from APSE phages associated with *Hamiltonella defensa* in aphids and whiteflies. Preliminary screening for homologs of these genes in the scale insect symbionts was performed using OrthoFinder v2.5.4^148^, which identified putative orthologs across our bacterial genome dataset. To refine this analysis, we constructed custom BLAST databases containing the retrieved genes involved in reproductive manipulation and defensive symbiosis. Protein-coding genes predicted from the genomes of scale insect symbionts were queried against these databases using BLASTp (E-value ≤ 1e-5, minimum identity ≥ 30%). Candidate gene hits were further validated through phylogenetic analyses as described above. Protein sequences of candidate genes were aligned with homologous sequences retrieved from databases.

**Sup Fig. 1.**
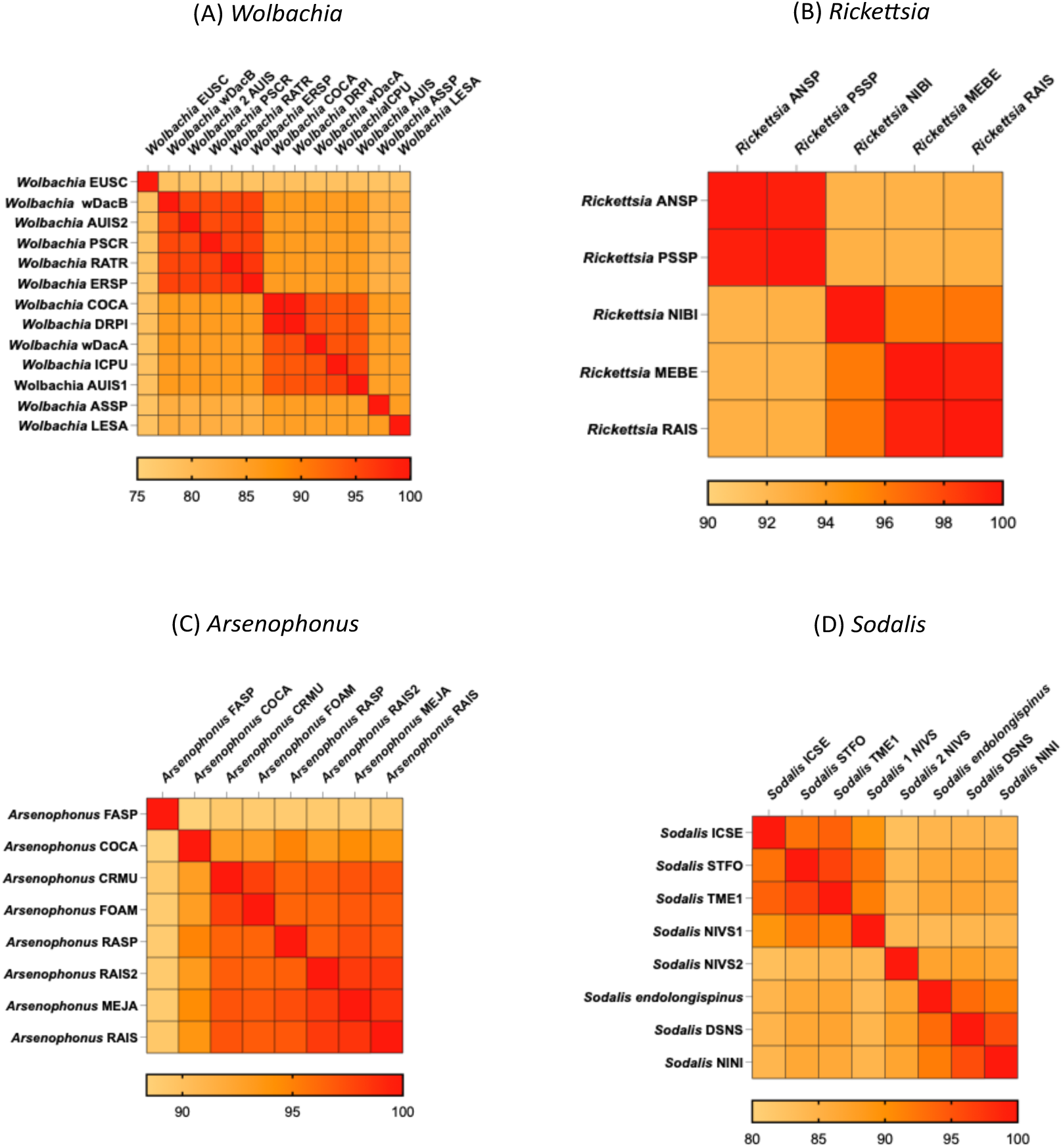
Genomic divergence among strains of four genera of bacterial symbionts in scale insects. Average Nucleotide Identity heatmaps of *Wolbachia* (A), *Rickettsia* (B), *Arsenophonus* (C), and *Sodalis* (D).

**Sup Fig. 2.**
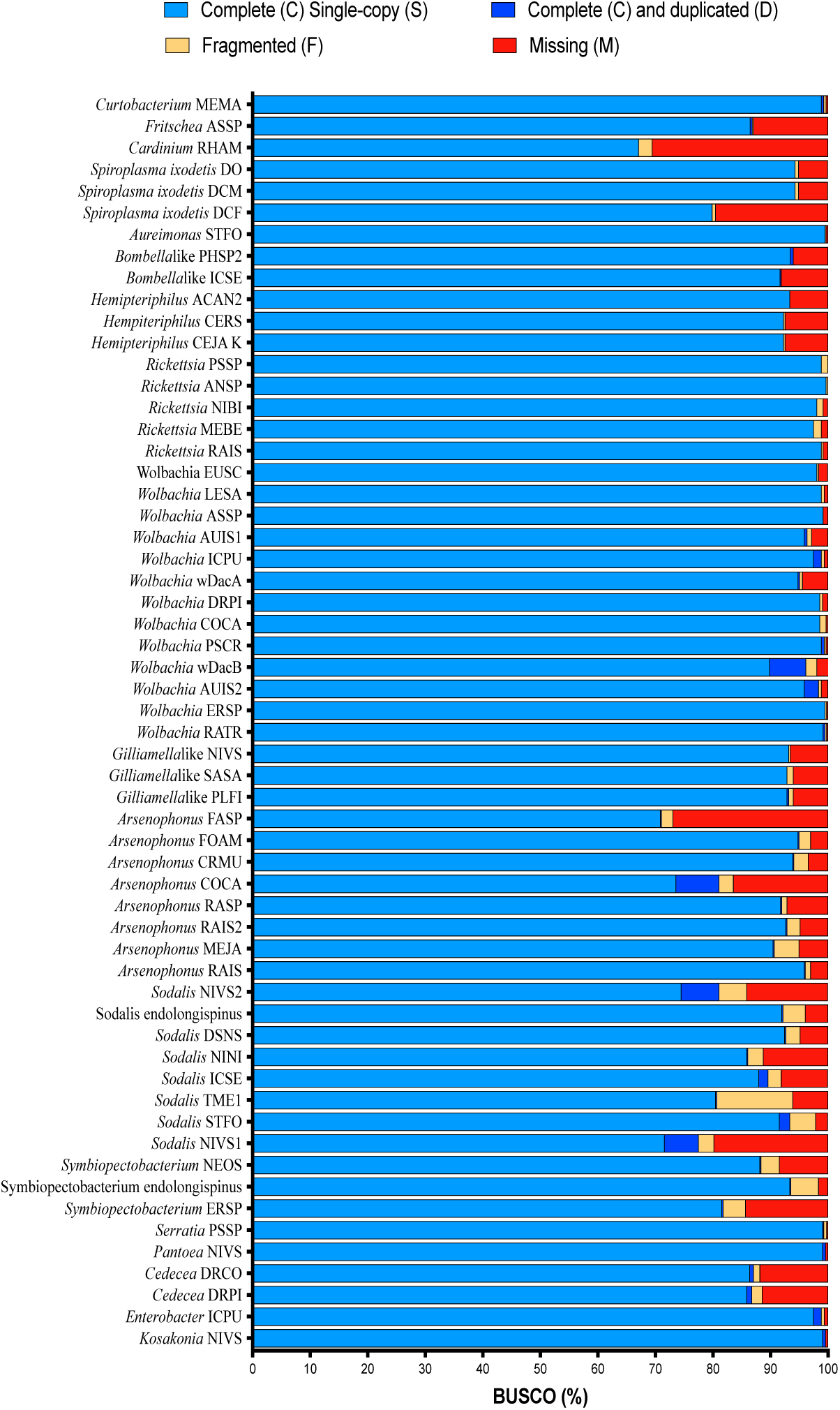
Assessment of genome completeness for scale insect symbionts using BUSCO scores. The horizontal bar plot represents the BUSCO scores (%) for each bacterial genome analyzed. Each bar is divided into four categories: Complete (C), Single-Copy (S) genes (light blue), Complete (C) and Duplicated (D) genes (dark blue), Fragmented genes (yellow), and Missing genes (red). The proportions of each category represent the completeness, redundancy, and quality of the bacterial genome assemblies.

**Sup Fig. S3.**
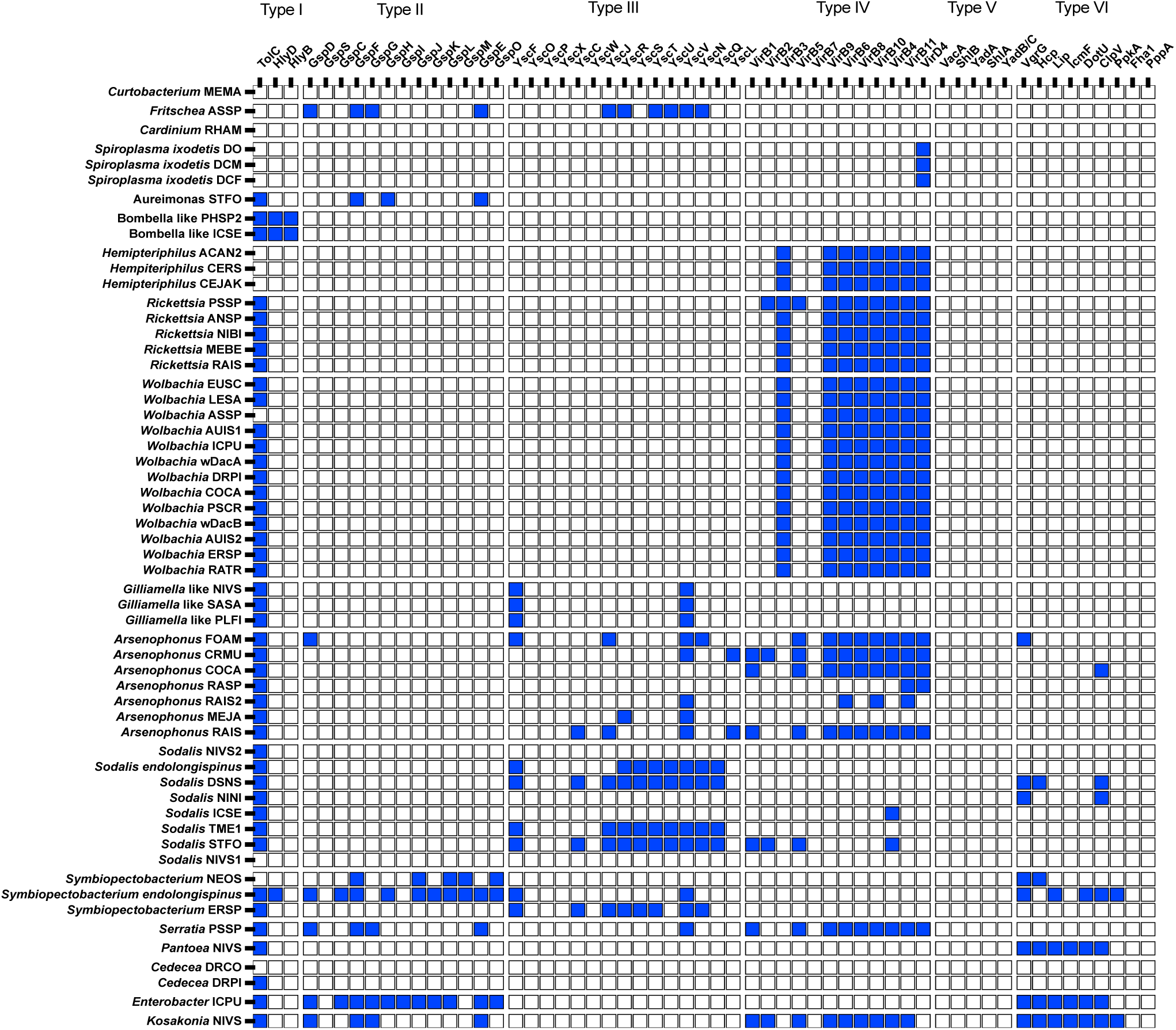
Comparative analysis of bacterial secretion systems across the symbionts from scale insects. Core components of the type I, type II, type III, type IV, type V, and type VI systems (as predicted by GhostKOALA) are shown. The presence of genes is indicated by squares filled in blue. Absence of genes is indicated by squares filled in white.

**Sup Fig. S4.**
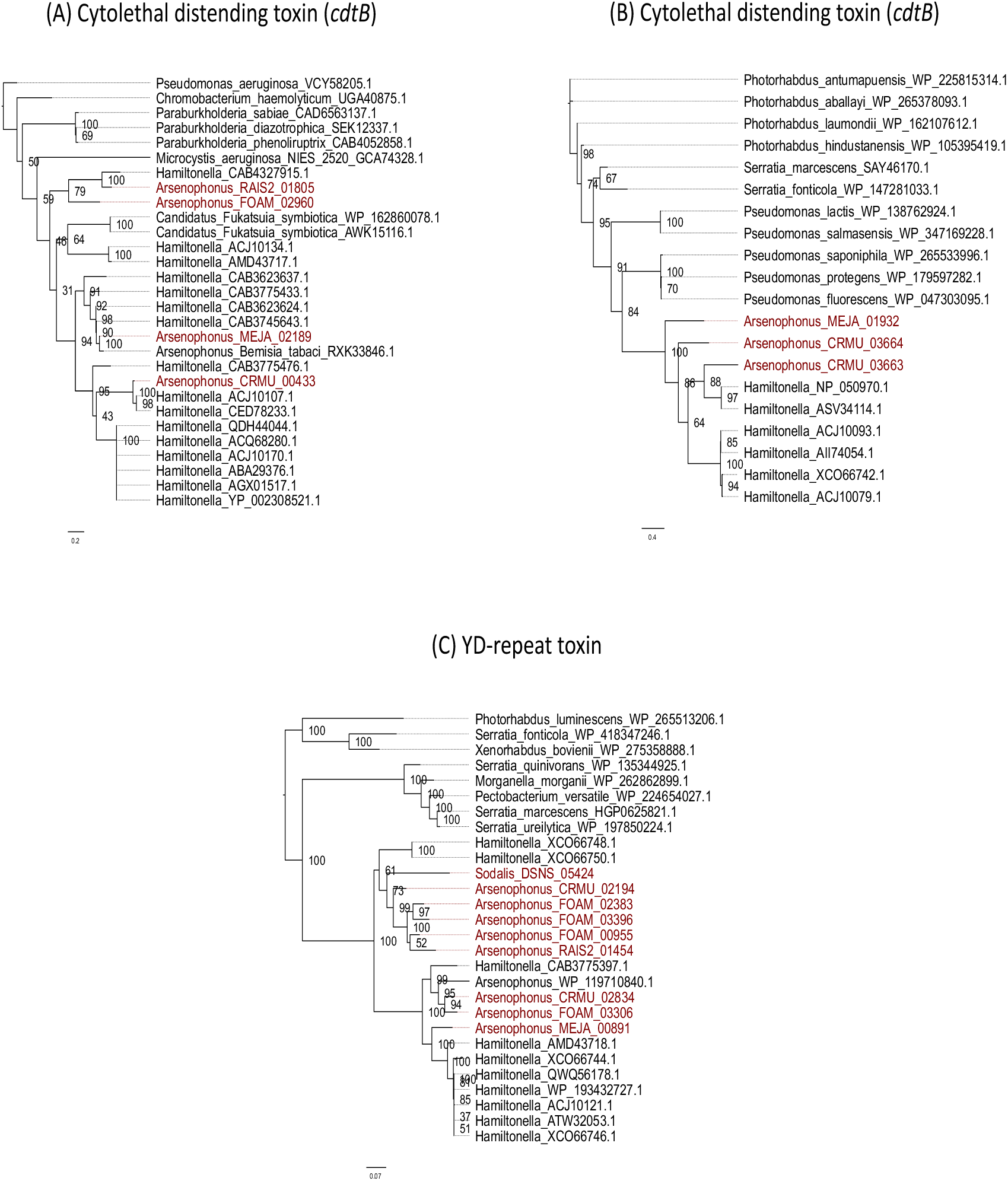
**Maximum likelihood tree** generated using IQtree based on nearly complete sequences of APSE phage encoded (A) cytolethal distending toxin (*cdtB*), (B) Shiga-like toxin (*stxA*), (C) YD-repeat toxins. Sequences from bacterial symbionts from scale insects and other insects, along with respective putative toxins from free-living and pathogenic bacteria are included.

**Sup Fig. S5.**
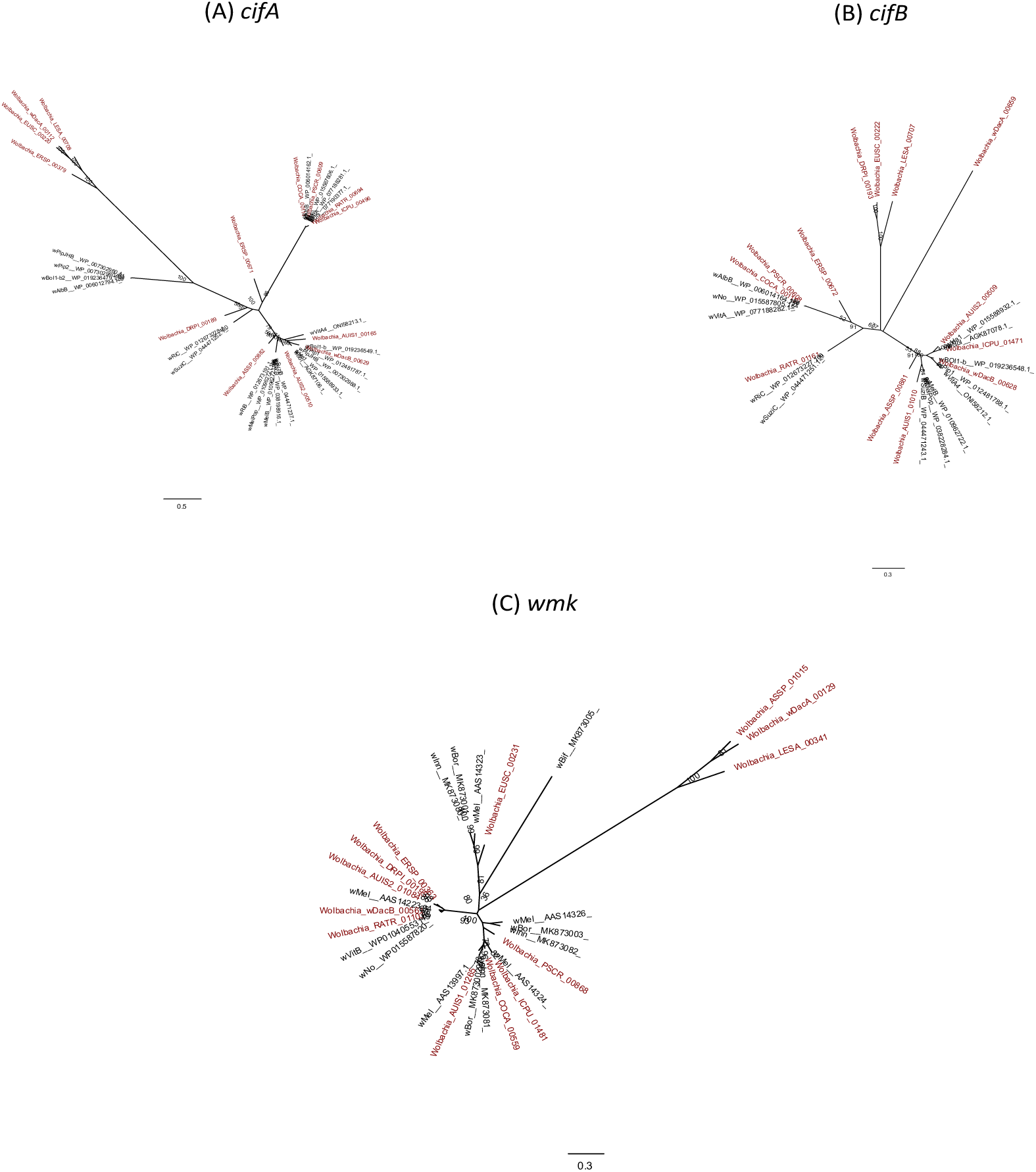
**Maximum likelihood tree** generated using IQtree based on nearly complete sequences of reproductive manipulation-related genes (A) *cifA*, (B) *cifB*, and (C) *wmk* of *Wolbachia* symbionts from scale insects, other insects and nematodes.

